# Precise transcriptional control of cellular quiescence by BRAVO/WOX5 complex in Arabidopsis roots

**DOI:** 10.1101/2020.07.15.203638

**Authors:** Isabel Betegón-Putze, Josep Mercadal, Nadja Bosch, Ainoa Planas-Riverola, Mar Marquès-Bueno, Josep Vilarrasa-Blasi, David Frigola, Rebecca Corinna Burkart, Cristina Martínez, Yvonne Stahl, Salomé Prat, Marta Ibañes, Ana I. Caño-Delgado

## Abstract

Root growth and development are essential features for plant survival and the preservation of terrestrial ecosystems. In the Arabidopsis primary root apex, stem-cell specific transcription factors BRAVO and WOX5 co-localize at the Quiescent Center (QC) cells, where they repress cell division so that these cells can act as a reservoir to replenish surrounding stem cells, yet their molecular connection remains unknown. Here, by using empirical evidence and mathematical modeling, we establish the precise regulatory and molecular interactions between BRAVO and WOX5. We found that BRAVO and WOX5 regulate each other besides forming a transcription factor complex in the QC necessary to preserve overall root growth and architecture. Our results unveil the importance of transcriptional regulatory circuits at the quiescent and stem cells to the control of organ initiation and growth of plant tissues.

## INTRODUCTION

Roots are indispensable organs to preserve plant life and terrestrial ecosystems under normal and adverse environmental conditions. In *Arabidopsis thaliana* (Arabidopsis), the primary root derives from the activity of the stem cells located at the base of the meristem in the root apex (Dolan et al, 1993; van den Berg et al, 1995). The root stem cell niche (SCN) is composed of a set of proliferative stem cells that surround the mitotically less active cells, named the quiescent centre (QC) (Scheres, 2007). Proximally to the QC, the vascular stem cells (VSC, also called vascular initial cells) give rise to functional procambial, xylem and phloem conductive vessels in the plant (De Rybel et al, 2016). Distally to the QC, the columella stem cells (CSC) give rise to the columella cells (Figure S1, (Gonzalez-Garcia et al, 2011; Stahl et al, 2009). The QC prevents differentiation of the surrounding stem cells (van den Berg et al, 1997), and its low proliferation rate provides a way to preserve the genome from replication errors. It also acts as a root stem cells reservoir, having the ability of promoting its own division rate to replenish the stem cells when they are damaged (Fulcher & Sablowski, 2009; Lozano-Elena et al, 2018).

BRASSINOSTEROIDS AT VASCULAR AND ORGANIZING CENTER (BRAVO) and WUSCHEL RELATED HOMEOBOX 5 (WOX5) are two transcription factors that are expressed in the QC and control its quiescence, as mutation of either BRAVO or WOX5 promotes QC cell division (Forzani et al, 2014; Pi et al, 2015; Vilarrasa-Blasi et al, 2014). BRAVO is an R2R3-MYB transcription factor and besides being expressed at the QC is also present at the vascular initials (Vilarrasa-Blasi et al, 2014). It was identified as a target of Brassinosteroid (BR) signaling, being directly repressed by BRI1-EMS-SUPPRESSOR 1 (BES1), one of the main effectors of the BR signaling pathway, altogether with its co-repressor TOPLESS (TPL) (Espinosa-Ruiz et al, 2017; Vilarrasa-Blasi et al, 2014). WOX5 is a member of the WUSCHEL homeodomain transcription factor family and it is localized mainly at the QC and to a lesser extent at the surrounding CSC and vascular initials (Pi et al, 2015; Sarkar et al, 2007). WOX5 can repress QC divisions by repressing CYCLIN D3;3 (Forzani et al, 2014), and in contrast with BRAVO, is also involved in CSC differentiation, as in the *wox5* mutant CSC differentiate prematurely (Sarkar et al, 2007).

Although BRAVO and WOX5 are well-studied plant cell-specific repressors of QC division, their molecular connection and the biological relevance in SCN proper functioning has not yet been established. In this study, we set the regulatory and molecular interactions between BRAVO and WOX5 at the SCN and disclose a common role as regulators of primary and lateral root growth and development. Our results show that BRAVO and WOX5 promote each other expressions and can directly bind to form a protein regulatory complex. BRAVO/WOX5 protein interaction underlies their functions as QC repressors to maintain stem cell development, that is essential for root growth and adaptation to the environment.

## RESULTS

### BRAVO and WOX5 control QC division and lateral root density

We have previously shown that *bravo* mutants have a phenotype of increased divisions at the QC compared to the wild-type (WT) (Vilarrasa-Blasi et al, 2014) (Figure 1A, B), which resembles the one described for *wox5* mutants (Bennett et al, 2014; Forzani et al, 2014; Sarkar et al, 2007) (Figure 1C). To address BRAVO and WOX5 interplay at repressing QC divisions, we generated the double *bravo wox5* mutants (Materials and Methods, Table S1). The double *bravo wox5* background also exhibited increased cell division compared to the WT (Figure 1A, D). Importantly, the frequency of divided QC was similar to that of *bravo* and *wox5* single mutants (Figure 1E). The mutual epistatic effect of these mutations suggests that BRAVO and WOX5 function interdependently at the WT primary root apex to supress QC divisions.

**Figure 1:**
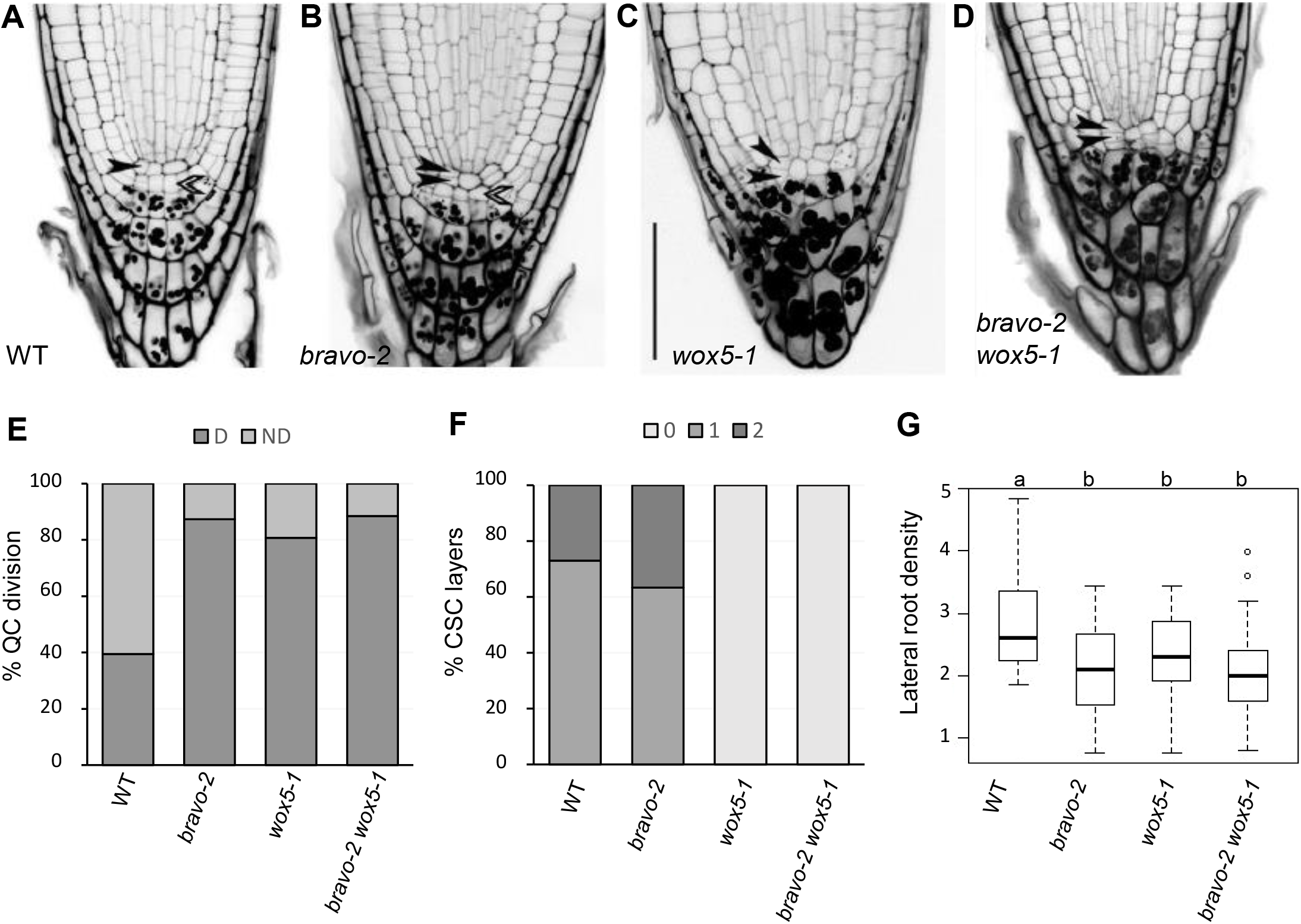
BRAVO and WOX5 are required for the QC identity and stem cells maintenance. **A-D)** Confocal images of mPS-PI stained 6-day-old seedlings of Col-0 (A), *bravo-2* (B), *wox5-1* (C) and *bravo-2 wox5-1* (D) mutants. Left black arrows indicate QC cells and right white arrows indicate CSC. Scale bar: 50 μm. **E)** Quantification of the QC divisions in 6–day-old roots expressed in percentage (n>50, 3 replicates). D: QC divided; ND: QC non divided. **F)** Quantification of CSC layers in 6-day-old roots expressed in percentage (n>50, 3 replicates). **G)** Lateral root density (number of lateral roots per mm of root length) of 7-day-olf WT, *bravo-2*, *wox5-1* and *bravo-2 wox5-1* mutants (n>40, 2 replicates). Different letters indicate statistically significant differences (p-value < 0.05 Student’s t-test).

Previous studies proposed that WOX5 represses CSC differentiation in a non-cell autonomous manner (Bennett et al, 2014; Sarkar et al, 2007), whereas no link was reported between this process and BRAVO, since the *bravo* mutants are not defective in CSC differentiation (Figure 1A, B, F). Genetic analysis showed that *bravo wox5* mutants display the same CSC differentiation as *wox5* single mutant (Figure 1A, C, D, F), corroborating that BRAVO does not control CSC differentiation (Vilarrasa-Blasi et al, 2014).

To address whether these stem cell-specific defects account for overall alterations in root growth and development, root architecture was analyzed. The *bravo wox5* double mutant shows slightly but significantly shorter roots than the WT (Figure S2A) and fewer lateral root density (Figure 1G). In the case of the lateral root density, 7-day-old *bravo wox5* seedlings show the same phenotype as the single mutants (Figure 1G), in agreement with previous reports for *wox5* (Tian et al, 2014a). Root growth defects become more exaggerated in the *bravo wox5* double mutant in 10-day-old seedlings (Figure S2B), therefore supporting the joint contributions of these two transcription factors to overall root growth and architecture.

### *BRAVO* and *WOX5* reinforce each other at the root stem cell niche

The QC division phenotype of the double *bravo wox5* mutant suggests an interplay between BRAVO and WOX5 at regulating QC divisions. Such interplay may take place through cross-regulation of their expressions. Indeed, we have previously shown that *WOX5* expression is reduced in the *bravo* mutant (Vilarrasa-Blasi et al, 2014), indicating that BRAVO regulates *WOX5* expression. To gain insight on the mutual regulatory activity of these two transcription factors, we thoroughly investigated *BRAVO* and *WOX5* expressions at the SCN in the single mutant and in the double *bravo wox5* mutant backgrounds.

In the WT primary root, *BRAVO* expression, reported by the *pBRAVO:GFP* line, is specifically located in the QC and the vascular initials (Vilarrasa-Blasi et al, 2014) (Figure 2A). The *pBRAVO* signal was increased in the *bravo* mutant (Figure 2B, H), suggesting that BRAVO negatively regulates its own expression. In contrast, in the *wox5* mutant*, pBRAVO* expression was strongly reduced, suggesting that WOX5 promotes *BRAVO* expression (Figures 2C, H). Inducible expression of WOX5 under the 35S promoter (35S:WOX5-GR) resulted in an increased *BRAVO* expression, as measured by RT-qPCR of root tips (Figure S3A). The fact that the increase is not as strong as the fold-induction of WOX5, suggests that WOX5 induces *BRAVO* only within the BRAVO native domain. Together, these results support that WOX5 activates *BRAVO* expression. Moreover, *pBRAVO* expression was equally reduced in the double *bravo wox5* mutant (Figure S4), as in the *wox5* mutant (Figure 2C, H), suggesting that BRAVO regulates its own expression aside the induction by WOX5. In the primary root, *WOX5* expression, as reported by the *pWOX5:GFP* line, is known to be mainly restricted to the QC, yet some expression is detected in the vascular initials (Pi et al, 2015) (Figure 2D). We found that *bravo* mutant displayed a significant reduction of *WOX5* expression (Figure 2E, I), supporting that BRAVO in turn induces expression of the *WOX5* gene. Further analysis of *WOX5* expression upon overexpressing *BRAVO* under an inducible 35S promoter (35S:BRAVO-Ei) showed that when BRAVO levels were induced, *pWOX5* levels remained similar to the WT, indicating that BRAVO is not able in its own to induce *WOX5* (Figure S3C-G). Together, these results support that BRAVO is necessary to maintain proper *WOX5* levels in the QC but does not induce them. Subsequently, an increased *pWOX5:GFP* expression towards the provascular cells was observed in the *bravo wox5* double mutant (Figure 2G), similar to *wox5* mutant (Figure 2F, I). These findings suggest that *WOX5* restricts its own expression to the QC, while BRAVO-dependent activation of *WOX5* acts upstream such *WOX5* autoregulation.

**Figure 2:**
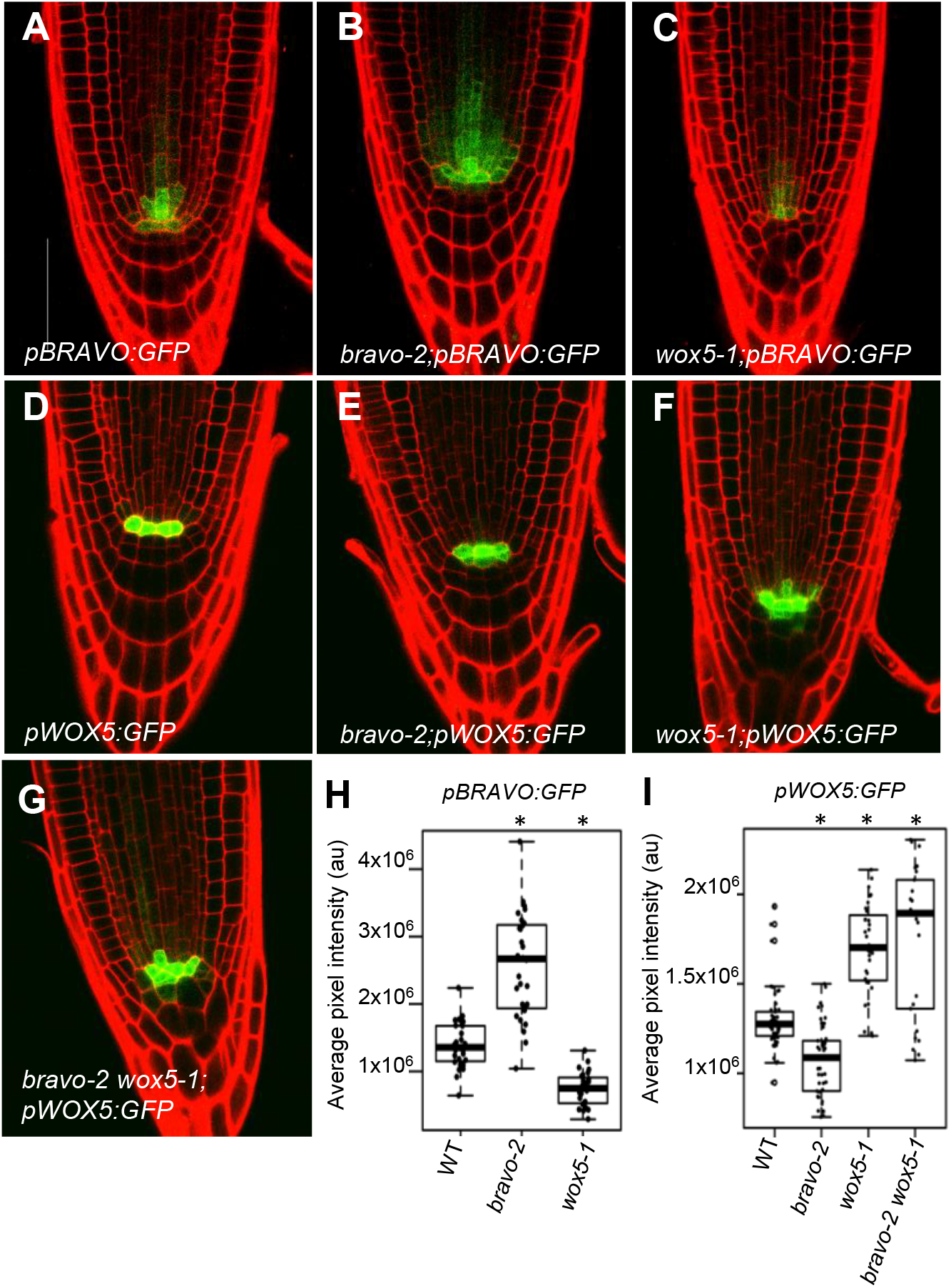
BRAVO and WOX5 reinforce each other at the root stem cell niche. **A-G)** Confocal images of PI-stained 6-day-old roots. GFP-tagged expression is shown in green. A-C) *pBRAVO:GFP* in WT (A), *bravo-2* (B) and *wox5-1* (C) knockout backgrounds. D-G) *pWOX5:GFP* in the WT (D), *bravo-2* (E), *wox5-1* (F) and *bravo-2 wox5-1* (G) knockout backgrounds. Scale bar: 50 μm. **H, I)** Quantification of the GFP fluorescent signal of the roots in A-C (H) and D-G (I). Boxplot indicating the average pixel intensity of the GFP in the stem cell niche. (n>25, 3 biological replicates, *p-value < 0.05 Student’s *t*-test for each genotype versus the WT in the same condition).

Brassinolide (BL) is the most active BR hormone compound. BL treatment is known to modify *BRAVO* and *WOX5* expression, by reducing the first and increasing the second of these genes (Gonzalez-Garcia et al, 2011; Vilarrasa-Blasi et al, 2014) (Figures S4 and S5). We found that when roots were grown on BL, the changes in *BRAVO* and *WOX5* expressions in *bravo*, *wox5* and the *bravo wox5* double mutant respect to the WT exhibited the same trends as when plants were grown in control media without BL (Figures S4 and S5). These results suggest that the mutual regulation of BRAVO and WOX5, as well as their autoregulation, is not significantly altered by BL treatment.

### *WOX5* induces *BRAVO*, which alleviates *WOX5* self-inhibition

To provide a comprehensive scheme of *BRAVO* and *WOX5* cross-regulation in the SCN able to account for the changes in expression levels observed in the various mutant backgrounds, we turned into mathematical modeling (Material and Methods). Because *BRAVO* is induced in the WOX5 overexpression line (Figure S3A) and *BRAVO* expression decreases in the *wox5* mutant (Figure 2C), the model considered that WOX5 induces (either directly and/or through intermediate molecules) the expression of *BRAVO* (Figure 3A). To account for the increase in *pBRAVO* expression in the *bravo* background (Figure 2B), the model assumed that BRAVO drives an effective inhibition on its own expression (Figure 3A), probably in an indirect manner. The model indicates that these two regulations can drive a decrease in *BRAVO* expression in the *bravo wox5* double mutant (Figure 3B), as found by the GFP expression data (Figure S4). Therefore, the model indicates that these two regulations on *BRAVO* are sufficient to account for its levels of expression in the single and double mutants (Figure 3B).

**Figure 3:**
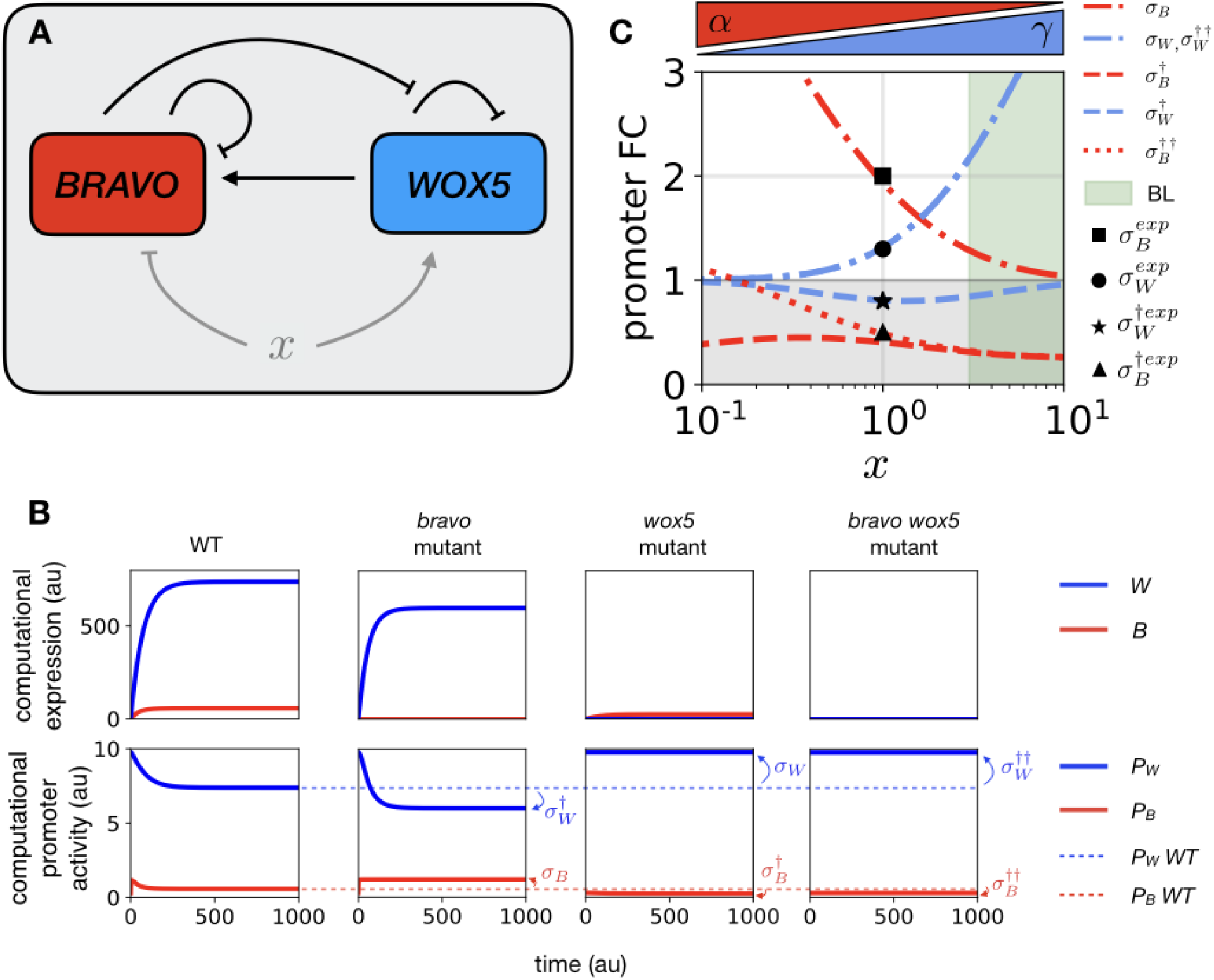
*WOX5* activates *BRAVO*, which in turn alleviates *WOX5* self-inhibition in the stem cell niche. **A)** Schematic representation of the effective regulations in the SCN between *BRAVO* and *WOX5*: *BRAVO* feeds back on its own activity by reducing it and is activated by *WOX5*. *WOX5* also feeds back on its own activity by reducing it, a regulation that becomes partially impaired by *BRAVO*. Additional factors *x* can be regulating both *BRAVO* and *WOX5* or either one. We exemplify one such a factor that regulates both, by downregulating *BRAVO* and upregulating WOX5. *x* can be understood as BR signaling. Arrows denote activation and bar-ended lines denote inhibition. **B)** Model solutions for the temporal evolution of expression and promoter activities for the WT and mutants using as initial condition all activities set to zero (*B*(t=0)=0,*W*(t=0)=0) and parameter values as in Table S1. This time-evolution does not intend to mimic any data but is only shown to depict the changes in the stationary levels between WT and each mutant. Manifest in the panels are the fold-changes in promoter activities in the mutant compared to the WT (σ) as defined in Material and Methods. **C)** Fold-changes in promoter activity (σ) in the mutant compared to the WT predicted by the mathematical model as a function of the control parameter *x*. This control parameter increases WOX5 and reduces BRAVO promoter activities (blue and red triangles; according to α=0.3/*x*, γ=250*x*/(*x*+9)). *x*=1 corresponds to the CTL condition, while *x*>1 can mimic BL conditions (green shaded area). The experimentally observed values in CTL conditions (computed as ratios of the median GFP) are drawn as black markers (see legend). The experimental fold-changes corresponding to the double mutants are not shown, as are assumed to be equal to the single mutants within the confidence interval of the experiments 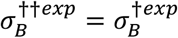 and 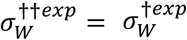. Error bars of these data (which can span ranges ±σ) are not depicted for clarity. In the plot, the region of fold change FC<1 (i.e. the promoter activity is reduced in the mutant) is shaded in gray to visually distinguish it from the region where FC>1 (i.e. the promoter activity is increased in the mutant).

Because *pWOX5* expression in the SCN increases in the *wox5* mutant (Figure 2F), the model considered that WOX5 represses (directly or indirectly) its own promoter activity (Figure 3A). In addition, the model assumed that BRAVO inhibits partially this repression (Figure 3A). With these regulations, the model accounts for the increase of *WOX5* expression in the *bravo* mutant, as well as for the *WOX5* decreased expression in the *wox5* and *bravo wox5* mutants (Figure 3B), as we found in the GFP expression studies (Figure 2F,G). Therefore, the model proposes that BRAVO promotes *WOX5* expression by alleviating *WOX5* self-inhibition.

With these interactions, the model precisely captures all changes in *BRAVO* and *WOX5* expression in the *bravo*, *wox5* and *bravo wox5* mutants (Figure 3B, C). In the model, parameter values were adjusted such that the fold-changes between promoter activities in the single mutants compared to the WT matched the fold-changes in GFP expressions of our empirical data (Figure 3C, Material and Methods). In addition, these values were restricted such that under control conditions *pBRAVO* expression is lower than *pWOX5* expression in the WT (Figure 3B), as suggested by GFP expression (Material and Methods) and RNAseq of the root tip (Clark et al, 2019).

The model indicates that the trends in the changes of expression levels between each mutant and the WT are maintained when the rate of BRAVO promoter activity decreases and/or the rate of WOX5 promoter activity is increased (Figure 3C). This is in agreement with the results obtained upon BL treatment (Figure S4 and S5), which reduces *BRAVO* expression whereas it increases *WOX5* expression.

### BRAVO and WOX5 directly interact into a transcriptional complex

Our results so far support that BRAVO and WOX5 reinforce each other at the SCN. To further decipher BRAVO and WOX5 interplay, we next evaluated the possible physical interaction between the BRAVO and WOX5 proteins. Using Förster resonance energy transfer measured by fluorescence lifetime microscopy (FRET-FLIM) (Figure 4A-K) and yeast two-hybrid assays (Figures 4L and S6A) we observed that BRAVO can directly interact with WOX5 (Figure 4B, G, K and L), which indicates that BRAVO and WOX5 form a transcriptional complex.

**Figure 4:**
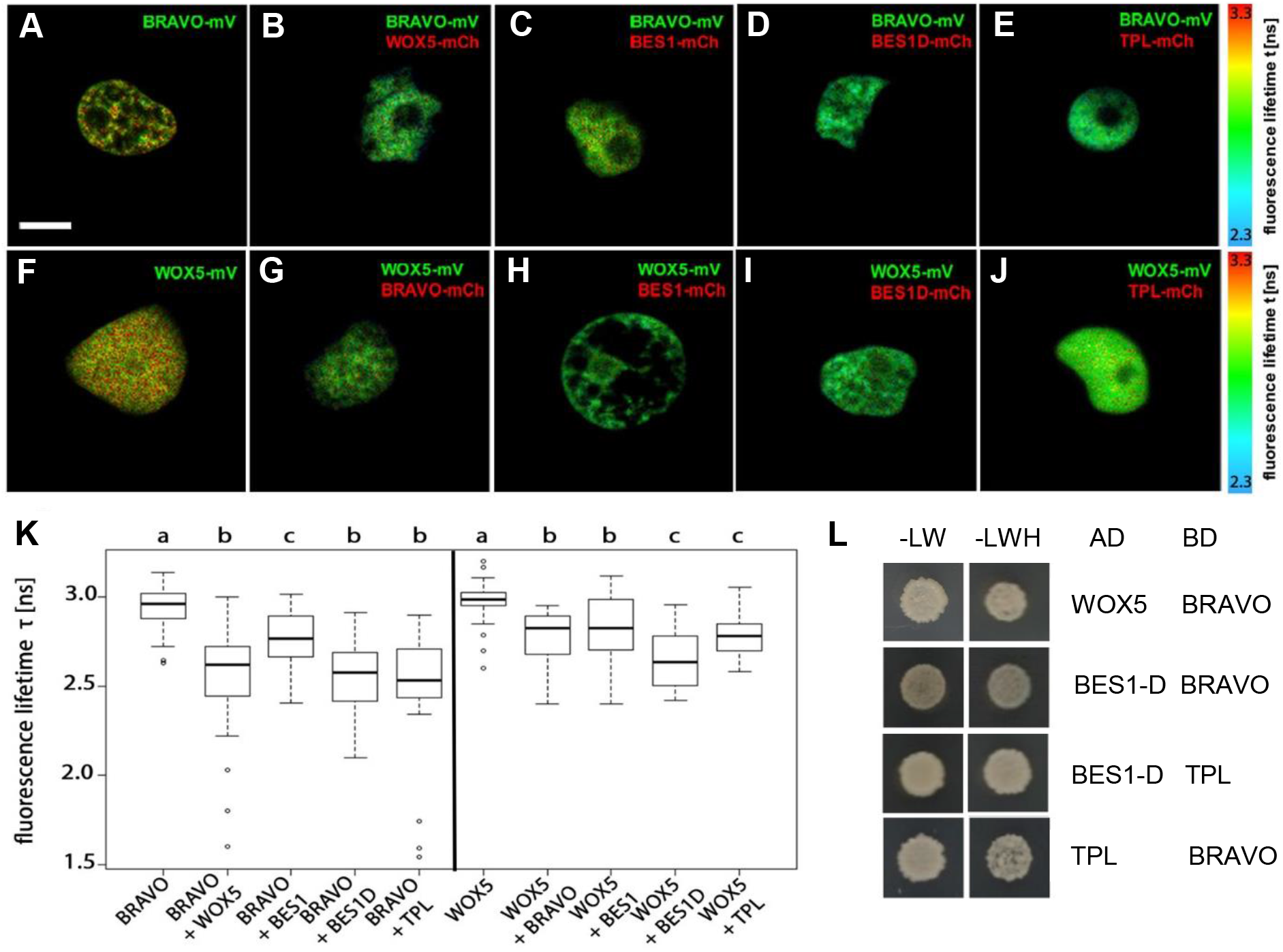
BRAVO interacts with WOX5. **A-J)** Interaction of BRAVO with WOX5 (B), BES1 (C), BES1-D (D) and TPL (E); and interaction of WOX5 with BRAVO (G), BES1 (H), BES1-D (I) and TPL (J) measured by FRET-FLIM. GFP fluorescence lifetime τ [ns] was measured in transiently expressing *Nicotiana benthamiana* leaf epidermal cells. GFP fluorescence lifetime fitted pixel-wise with a mono-exponential model of BRAVO and WOX5 interactions. mV, mVenus; mCh, mCherry. Scale bar: 5 μm. **K**) Fluorescence-weighted average lifetimes of BRAVO and WOX5 interactions fitted with a double-exponential model of the indicated samples are summarized in box plots. Statistical significance was tested by one-way ANOVA with a Sidakholm post-hoc test. Different letters indicate statistically significant differences (p<0.01; n>20). **L)** Yeast two-hybrid assay showing BRAVO interacting with WOX5, BES1-D and TPL. In the left column yeast cells were grown on control media, and in the right column yeast cells were grown on control media lacking Leu, Trp and His, indicating an interaction between the proteins.

As we previously demonstrated that the BR-regulated BES1/TPL complex acts as a transcriptional repressor of BRAVO transcription (Espinosa-Ruiz et al, 2017; Vilarrasa-Blasi et al, 2014), in addition to BES1 directly interact with BRAVO (Vilarrasa-Blasi et al, 2014), and TPL is shown to interact with WOX5 (Pi et al, 2015), we further investigated binding of BRAVO and WOX5 to these transcriptional regulators. We found that both BRAVO and WOX5 physically interact with BES1, and this interaction was stronger for the active BES1-D protein (Yin et al, 2002) (Figures 4C, D, H, I, K), consistent with our previous findings that the of BES1 EAR domain is necessary for BES1/BRAVO interaction (Vilarrasa-Blasi et al, 2014); Figure S6A). Our analysis shows that BES1 binds to WOX5 (Figures 4H, I, K and S6C) with an equivalent affinity as to BRAVO (Figures 4K and S6B), and that this interaction is stronger with BES1-D (Figure 4K). Moreover, both BRAVO and WOX5 were also observed to interact with the co-repressor TPL (Figures 4E, J, K, L and S6). Collectively, these data show that BRAVO and WOX5 directly interact to form a transcriptional complex, and that each can bind active BES1 and TPL, suggesting these proteins are able to compete for their mutual binding.

### BRAVO-WOX5 complex is relevant for the control of QC divisions

The equal divided QCs in the double *bravo wox5* mutant compared to the single mutants (Figure 1A-E) suggests that BRAVO and WOX5 interplay at repressing QC divisions. We found two ways for this interplay to take place: through mutual regulation of their expressions (Figures 2, 3A) and through the formation of a protein BRAVO-WOX5 complex (Figure 4A-K). We turned into mathematical modeling to assess the contribution of each of these regulations to the phenotype of divided QCs (Material and Methods). We set a regulatory function for the frequency of divided QCs that explicitly incorporates the individual contributions mediated by BRAVO (TB) and by WOX5 (TW) and the jointly mediated contribution by both BRAVO and WOX5 together (hereafter named “joint contribution”, T_BW_) (Material and Methods). In this regulatory function, the joint contribution (TBW) is the one that takes into account the existence of the BRAVO-WOX5 complex. In contrast, the mutual regulations of *BRAVO* and *WOX5* expressions act independently from the joint contribution and are only included in the individual contributions (i.e. T_B_ and T_W_). Specifically, since *WOX5* expression decreases in the *bravo* mutant (Figure 2I), we reasoned that individual WOX5 repression of QC divisions is attenuated by a factor q_W_^Bm^<1 in the *bravo* mutant compared to the WT (Material and Methods). Similarly, to take into account the regulation that WOX5 makes on *BRAVO* expression, we considered that the individual contribution by BRAVO was attenuated by a factor q_B_^Wm^ in the *wox5* mutant compared to that in the WT (q_B_ ^Wm^ <1). Because the extent of these attenuations and hence the values of q_W_^Bm^ and q_B_^Wm^ (which range from 0 to 1) cannot be measured, we estimated them through the fold-changes in expression in the mutants as follows (Materials and Methods). We used q_W_^Bm^=0.8, which is similar to the fold-change of *WOX5* expression in the *bravo* mutant compared to the WT (Figures 2I, 3C). The fact that *wox5* exhibits phenotypes that are absent in the *bravo* mutant, such as CSC differentiation, also suggests that q_W_^Bm^ is not too small. The estimate for q_B_^Wm^ based on the fold-change of *BRAVO* expression in the *wox5* mutant is q_B_^Wm^=0.5 (Figures 2H, 3C). Yet, from the root phenotypes of the mutants we cannot exclude other, e.g. smaller, values. Therefore we evaluated the model results for different values of q_B_^Wm^.

We used the experimental data on the frequency of divided QCs in the WT, the single mutants and the double mutant (Figure 1E), with an estimation of their confidence intervals (Material and Methods), to extract which are the individual contributions (i.e. the BRAVO-mediated and the WOX5-mediated) as well as the joint BRAVO-WOX5 contributions in the WT (Material and Methods). For intermediate q_B_^Wm^ values (q_B_^Wm^ >0.4 upwards, being q_B_^Wm^=0.5 the estimate from fold-change *BRAVO* expression in the *wox5* mutant), the model results show that in the WT the joint contribution of BRAVO-WOX5 is the only one relevant (Figure 5A). Therefore, the analysis indicates that the joint BRAVO-WOX5 contribution is essential to describe the QC division data if BRAVO and WOX5 control each other action on QC division only partially. Individual BRAVO contribution becomes relevant only for small q_B_^Wm^ values, i.e. only if BRAVO’s role on QC division is mostly controlled by WOX5. Yet in this scenario, which would correspond to BRAVO acting downstream of WOX5 to repress QC divisions, the model indicates that the joint contribution of BRAVO and WOX5 is also relevant to the regulation of QC divisions in the WT, regardless of its specific activatory/inhibitory role (Figure 5A). Taken together, our analyses highlight the significant contribution of the BRAVO/WOX5 heterodimeric complex in the control of QC divisions, to the preservation of the normal growth and development of primary and lateral root organs in the plant.

**Figure 5:**
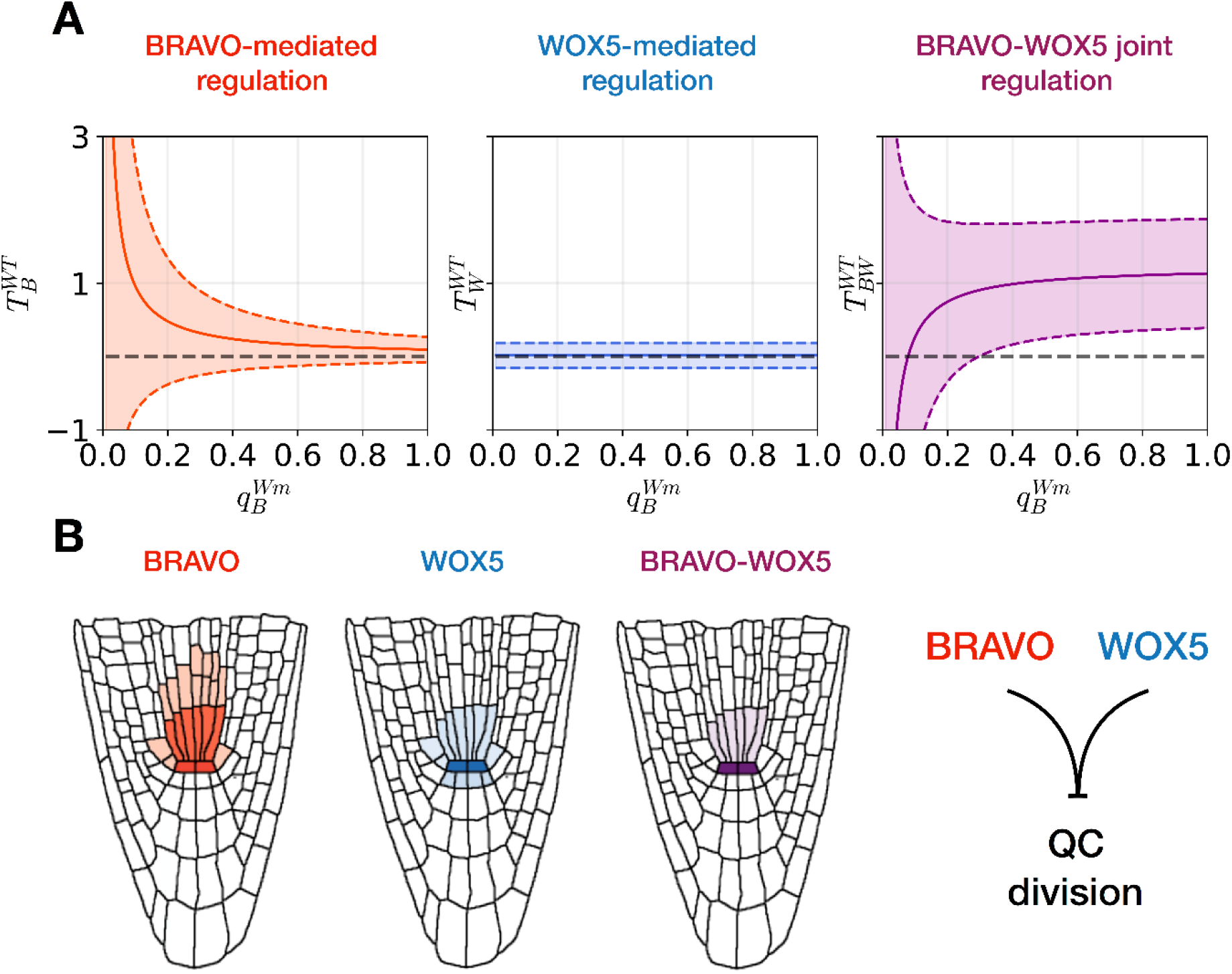
BRAVO and WOX5 have a joint role in repressing QC divisions. **A)** Computational estimation of the contributions of BRAVO-mediated 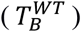, WOX5-mediated 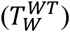 and BRAVO-WOX5 joint 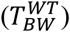 regulations of QC divisions in the WT, as a function of the attenuating factor of BRAVO contribution in the *wox5* mutant, 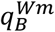. Continuous lines represent the best estimated values, while dashed lines are the enveloping confidence intervals (e.g. 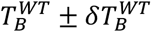). The horizontal grey dashed lines mark the zero lines. For a wide range of 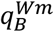 values, the joint contribution of BRAVO and WOX5 is important, while the individual contribution of BRAVO only increases for small values of 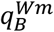. In all three panels, we set 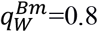. Positive contributions correspond to repression of QC divisions, while negative contributions correspond to activation of QC divisions. **B)** Sketch representing the spatial distribution of BRAVO, WOX5 and their product BRAVO x WOX5, which can be interpreted as the protein complex. Their joint interaction peaks at the QC, where repression of cell division occurs.

## DISCUSSION

In the Arabidopsis primary root, BRAVO and WOX5 are two transcription factors that repress QC divisions and whose expressions co-localize mostly at the QC (Forzani et al, 2014; Vilarrasa-Blasi et al, 2014). Our results show that BRAVO and WOX5 interplay at different levels to repress QC divisions. In addition, we show that the joint action of these cell-specific transcription factors promotes overall root growth and development.

Our data indicate that BRAVO and WOX5 mutually promote each other’s expressions. Hence, neither of them is downstream the other, yet their mutual regulations are very distinct. While WOX5 is able to induce *BRAVO*, BRAVO does not directly induce *WOX5* expression but it drives partial inhibition of *WOX5* self-regulation. These different regulatory mechanisms and the quantitative changes in gene expression they drive, suggest that the effect WOX5 on *BRAVO* and thereby on BRAVO-mediated regulation can be more relevant than the effect BRAVO has upon *WOX5* and WOX5-mediated action. This is consistent with the known SCN phenotypes of *bravo* and *wox5* mutants (Bennett et al, 2014; Forzani et al, 2014; Pi et al, 2015; Sarkar et al, 2007; Vilarrasa-Blasi et al, 2014), where *wox5* exhibits, besides a similar increased QC division phenotype as *bravo*, an overall distorted and disorganized SCN morphology and CSC premature differentiation that is absent in the *bravo* mutant.

The mutual regulation between BRAVO and WOX5 involves WOX5 inhibition of its own expression while it induces that of *BRAVO*, which in turn reverses WOX5 self-repression. Based on our data, it can be suggested that *WOX5* self-inhibition is through WOX5 bound to TPL and that BRAVO attenuates it by competing with TPL for binding WOX5. Moreover, BRAVO is found to ultimately down-regulate its own expression, although this probably occurs through other intermediate molecules, as BRAVO has been shown to activate itself by directly binding its own promoter (Vilarrasa-Blasi et al, 2014). By evaluating expression changes between the WT and the mutants we gained information on the overall BRAVO-WOX5 regulatory system. Its regulation results from the direct binding of these proteins to their promoters and from the transcriptional control driven by them, as far as these proteins bind each other and to additional regulators. Hence, interactions here described are effective in the sense that they are the result of multiple, direct and indirect, regulatory mechanisms. For instance, *WOX5* self-repression can also involve a negative feedback where WOX5 activates a repressor or represses an activator, among other possibilities. In this context, control of auxin-ARF and auxin-IAA (Tian et al, 2014b) as well as the PLETHORA genes (Burkart et al, 2019) were all shown to involve negative feedbacks with *WOX5*. WOX5 induction of *BRAVO* expression could be as well through a downstream target of WOX5.

Another important molecular link between BRAVO and WOX5 as revealed by our data is their physical protein-protein interaction. The QC is where these two transcription factors mostly co-localize, which suggests that they act as co-partners of a single complex only in the QC, where they converge. The consistent and overlapping role of BRAVO and WOX5 at promoting lateral root development also points to a relevant role of the BRAVO-WOX5 complex for this function.

Our analysis supports that QC division is controlled via BRAVO-WOX5 joint regulation, besides an additional regulation individually mediated by BRAVO. This joint regulation is expected to be mediated by BRAVO-WOX5 physical interaction. This scenario explains the phenotype of increased divisions at the QC upon BL treatment (Gonzalez-Garcia et al, 2011), by the response of BRAVO and WOX5 to this treatment and their respective roles as repressors of QC divisions. Actually, although the intensity and domain of expression of WOX5 increases in roots grown in BL medium, at the same time the BL treatment strongly represses BRAVO (Vilarrasa-Blasi et al, 2014). Hence, in the absence of its partner BRAVO, WOX5 no longer represses QC divisions in roots grown on BL. At a mechanistic level, the BRAVO-WOX5 protein complex may bind CYCLIN-D3:3, as shown to occur for WOX5 (Forzani et al, 2014).

Interestingly, we also found that BRAVO and WOX5 promote root growth and lateral root development. In LR development, the formation of the organizing center and the stem cell niche occurs after LR initiation (Banda et al, 2019). A high number of genes are commonly expressed at the SCN of primary and lateral roots, such as PLT, SHR, SCR or TCP (Goh et al, 2016; Shimotohno et al, 2018). Loss-of function of these genes leads to an increased number of aberrant lateral roots and reduced levels of *WOX5* (Shimotohno et al, 2018), and thus it is possible that BRAVO/WOX5 complex not only controls stem cell niche maintenance in the primary root, but also in the lateral roots.

Finally, our study sets a framework for future studies on the interplay between WOX5 and BR signaling in the control of CSC differentiation. WOX5 is known to repress CSC differentiation (Pi et al, 2015; Sarkar et al, 2007). However, upon BL treatment, and in *bes1-D* gain of function mutants, CSC differentiate prematurely (Gonzalez-Garcia et al, 2011), in apparent contradiction with the inhibitory role associated with WOX5, and its induced expression in these roots. One option comes from assuming that BL-induced CSC differentiation is independent from WOX5 and overrides WOX5-mediated repression. In this case, a tug-of-war between WOX5-mediated repression and BL-dependent activation of CSC differentiation would tip the balance in favor of BR-action. Another possibility is that BR downstream effectors such as BES1-D inactivate WOX5 and/or impede its function. An increase of BES1-D by BL may boost WOX5 sequestration into WOX5-BES1-D complexes, since we showed that WOX5 and BES1-D physically interact. Assuming these complexes inactivate WOX5 function, CSC differentiation would no longer be repressed by WOX5 in the presence of BL. Moreover, the fact that BES1-D directly interacts with TOPLESS, and this co-repressor also recruited by WOX5 to the inhibition of CSC differentiation (Pi et al, 2015), suggest that in plants treated with BL WOX5 function may further impaired by most of TPL being bound to BES1-D.

To conclude, understanding of signaling networks operating in stem cell development is becoming essential to decipher plant growth and adaptation to the environment. Systems biology approaches provide a closer picture to reality unveiling how complex and dynamics network of cell-specific transcription factors act to preserve stem cell function in plants. Here, untapping the action of two main regulators of quiescent cell division, BRAVO and WOX5, not only discloses that these factors operate as a transcriptional complex in preserving stem cell function, but also unveils their joint roles in primary and lateral root development.

## AUTHOR CONTRIBUTIONS

A.I.C-D. and M.I. designed and supervised the study. I.B-P., N.B., A.P-R, J.V-B. and M.M-B. performed the experiments. J.M., D.F. and M.I. performed the mathematical modeling. Y.S. and R.C.B. performed and analysed the FRET-FLIM assays. S.P. and C.M. collaborated in the Y2H and BiFC assays. I.B-P., J.M., N.B., M.I. and A.I.C-D. wrote the manuscript and all authors revised the manuscript.

## ACKNOWLEDGMENTS

A.I.C-D. is a recipient of a BIO2016-78955 grant from the Spanish Ministry of Economy and Competitiveness and a European Research Council, ERC Consolidator Grant (ERC-2015-CoG – 683163).

N.B. is funded by the FI-DGR 2016FI_B 00472 grant from the AGAUR, Generalitat de Catalunya; I.B-P. by the FPU15/02822 grant from the Spanish Ministry of Education, Culture and Sport; and A.P-R. by the SEV-2015-0533 from the Severo Ochoa Programme for Centers of Excellence in R&D. M.I. and J.M. acknowledge support from the Spanish Ministry of Economy and Competitiveness and FEDER (EU) through grant FIS2015-66503-C3-3-P, from Ministerio de Ciencia, Innovación y Universidades / Agencia Estatal de Investigación / Fondo Europeo de Desarrollo Regional, Unión Europea through grant PGC2018-101896-B-I00 and from the Generalitat de Catalunya through Grup de Recerca Consolidat 2014 SGR 878 and 2017 SGR 1061. J.M. is funded by the Spanish Ministry of Education through BES-2016-078218. Y.S. and R.C.D. are funded by the Deutsche Forschungsgesellschaft (DFG) (grant STA12/12 1-1). CRAG is funded by “Severo Ochoa Programme” from Centers of Excellence in R&D 2016-2019 (SEV-2015-485 0533).

## MATERIAL AND METHODS

### Plant Material and Root Measurement

All WT, mutants and transgenic lines are in the Arabidopsis ecotype Columbia (Col-0) background (Table S2). The double mutant *bravo wox5* was generated by crossing the *bravo* and *wox5* single mutants. The double mutant homozygous lines were selected by genotyping. The primers used for *bravo* and *wox5* genotyping are listed in Table S3.

Seeds were surface sterilized and stratified at 4°C for 48 hours before being plated onto 0.5X Murashige and Skoog (MS) salt mixture without sucrose and 0.8% plant agar, in the absence or presence of Brassinolide (Wako, Osaka, Japan). β-estradiol (30 μM) from Sigma diluted in DMSO was used to induce BRAVO expression for 6 days. Dexamethasone (1 μM) from Sigma diluted in EtOH was used to induce WOX5 expression for 6 days. For RT-qPCR experiments β-estradiol and dexamethasone treatments were applied for 24 hours.

Plates were incubated vertically at 22°C and 70% humidity in a 16 hours light/8 hours dark cycle. Primary root length was measured from plates images, using ImageJ (https://imagej.nih.gov/ij/) and MyROOT (Betegon-Putze et al, 2019) softwares. The lateral root density was calculated by dividing the total number of emerged lateral roots of individual seedlings by the mean of the root length of those seedlings.

### Confocal Microscopy and Quantification of Fluorescence Signal

Confocal images were taken with a FV 1000 Olympus confocal microscope after Propidium iodide (PI, 10 μg/ml) staining. PI and GFP were detected with a band-pass 570-670 nm filter and 500-545 nm filter, respectively. Images were taken in the middle plane of 6-day-old roots. The fluorescence intensity was quantified with ImageJ using the Integrated Density value obtained from individual plants. The quantified area was selected with a ROI that contained the SCN (Figure S6). The laser settings for *pBRAVO:GFP* and *pWOX5:GFP* are different, as WOX5 has a stronger expression than BRAVO. The analysis of *pBRAVO:GFP* in *bravo wox5* double mutant background was done with different confocal settings. The analysis of QC cell division and CSC differentiation was carried out by imaging fixed roots through a modified pseudoSchiff (mPS-PI) staining method (Truernit et al, 2008). Images were processed with the Olympus FV (Olympus, Tokio, Japan) and ImageJ software.

### RT-qPCR assay

RNA was extracted from root tip tissue with the Maxwell^®^ RSC Plant RNA Kit (Promega) using the Maxwell^®^ RSC instrument (Promega) according to the manufacturer’s recommendations, and concentrations were checked using NanoDrop 1000 Spectrophotometer (Thermo Fisher Scientific). cDNA was obtained from RNA samples by using the NZY First-Strand cDNA Synthesis Kit (NZYtech) according to the manufacturer’s recommendations. RT-qPCR amplifications were performed from 10 ng of cDNA using SYBR Green I master mix (Roche) in 96-well plates according to the manufacturer’s recommendations. The RT-qPCR was performed on a LightCycler 480 System (Roche). *ACTIN2* (AT3G18780) was used as housekeeping gene for relativizing expression. Primers used are described in Table S3.

### Yeast two-hybrid assay

Yeast two-hybrid assays were performed by the Matchmarker GAL4-based two-hybrid System (Clontech). Constructs were co-transformed into the yeast strain AH109 by the lithium acetate method (Gietz & Woods, 2002). The presence of the transgenes was confirmed by growth on SD-LW plates, and protein interaction was assessed by selection on SD-LWH plates. Interactions were observed after 4 days of incubation at 30°C.

### Transient expression in *Nicotiana benthamiana* for FLIM measurements

Preparation of transiently expressing *Nicotiana benthamiana* leaves and induction of fusion proteins tagged with either mVenus or mCherry by application of ß-estradiol was carried out as described in (Bleckmann et al, 2010).

### Acquisition of FLIM data

FLIM data acquisition was carried out using a confocal laser scanning microscope (LSM780 inverted microscope, Zeiss) equipped additionally with a time-correlated single-photon counting device with picosecond time resolution (Hydra Harp 400, PicoQuant). mVenus was excited at 485 nm with a pulsed (32 MHz) diode laser at 1.2 μW at the objective (40 x water immersion, C-Apochromat, NA 1.2, Zeiss). The emitted light was collected through the same objective and detected by SPAD detectors (PicoQuant) using a narrow range bandpass filter (534/35, AHF). Images were taken at 12.5 μs pixel time and a resolution of 138 nm/pixel in a 256×256 pixel image. A series of 40 frames was merged into one image and analysed using the Symphotime software package (PicoQuant).

### Analyses and presentation of FLIM data

The fluorescent lifetime of the collected photons in each merged image was analysed using the Symphotime software (PicoQuant). For this, a ROI covering the whole nucleus was created to reduce background fluorescence. All photons in this ROI were used to build a histogram of the fluorescence decay. A double-exponential fit model was used to approximate the intensity-weighted average fluorescence lifetime □ [ns] of all photons of the ROI. The instrument response function was measured with KI-quenched erythrosine and used for reconvolution in the fitting process (Weidtkamp-Peters & Stahl, 2017). The data from replicate measurements was summarized in box plots created in R software (https://www.r-project.org/). Statistical significance was tested by one-way ANOVA with a Sidakholm post-hoc test. Different letters indicate statistically significant differences (p < 0.01).

For the creation of FLIM images, photons from individual pixels of a merged image were analysed for fluorescent lifetime using the Symphotime software (PicoQuant). A mono-exponential fit model was used, as the photon number in each pixel was too low for a double-exponential model (Stahl et al, 2013). The individual pixels are colour-coded according to their fluorescence lifetime.

### Bimolecular fluorescence complementation assay (BiFC)

The *BRAVO* and *WOX5* coding sequences were inserted by LR-reaction (Invitrogen) into pBiFC binary vectors containing the N- and C-terminal YFP fragments (YFPN43 and YFPC43). Plasmids were transformed into the *Agrobacterium tumefaciens* GV3101 strain and appropriate combinations were infiltrated into *Nicotiana benthamiana* leaves (Occhialini et al, 2016). The p19 protein was used to suppress gene silencing. Infiltrated leaves were imaged two days after infiltration using an Olympus FV1000 laser scanning confocal microscope.

### Mathematical model of BRAVO and WOX5 effective regulations

We considered a model for the effective regulations that BRAVO and WOX5 perform on each other and on themselves in the SCN. In the model, *B* and *W* account for the total *BRAVO* and *WOX5* expression in the whole SCN. These expression levels are considered to be the product of the BRAVO and WOX5 promoter activities according to the following wild-type dynamics:

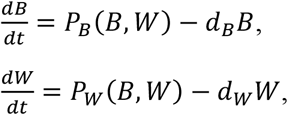

where *P_B_*(*B,W*) and *P_W_*(*B,W*) are the BRAVO and WOX5 promoter activities (production terms) respectively and *d_B_B* and *d_W_W* are the decay terms (assumed linear for simplicity, with decay rates *d_B_* and *d_W_*). To account for the regulation of the expression, each promoter activity depends on *BRAVO* and *WOX5* expressions. To compare with empirical data, we only considered the stationary state of the above dynamics (i.e. when time derivatives are equal to zero, 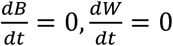). In the stationary state, *BRAVO* expression is proportional to BRAVO promoter activity (*B = P_B_*(*B,W*)/*d_B_*) and *WOX5* expression is proportional to WOX5 promoter activity (*W = P_W_*(*B,W*)/*d_W_*). Therefore, we used the promoter activity in the stationary state as the computational model read-out to be compared with the empirical data on *pBRAVO:GFP* and *pWOX5:GFP*.

Promoter activity terms *P_B_*(*B,W*) and *P_W_*(*B,W*) correspond to functions that describe the effective regulations that each expression ultimately performs on each promoter activity (see Figure 3A for a cartoon of these regulations). These effective regulations involve several intermediate steps, including translational and post-translational processes, and additional molecules. These are not explicitly modelled but are all together absorbed in the functionalities of *P_B_*(*B,W*) and *P_W_*(*B,W*). We expect these functions to be non-linear and we used continuous Hill-like functions exhibiting saturation with exponents larger than 1 (see parameter values in Table S1);

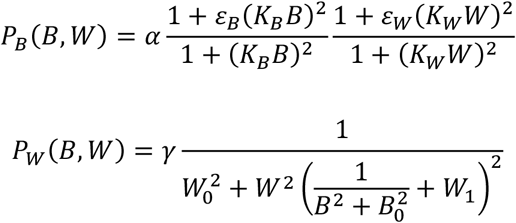

The BRAVO promoter activity *P_B_*(*B,W*) has: i) a basal production rate α, independent of *BRAVO* and *WOX5* expressions since our GFP data show that BRAVO promoter has activity in the double mutant *bravo wox5* (Figure S2). ii) A term that sets the activation of *BRAVO* expression by *WOX5*, with *WOX5* expression threshold value 1/K_W_ and activation strength εW > 1. According to this term, the production of *BRAVO* increases to αεW > α if *WOX5* expression is very high (*W*>>1/K_W_) and there is no *BRAVO*. iii) A term that accounts for the reduction of *BRAVO* expression by itself, with *BRAVO* expression threshold value 1/KB and inhibition strength εB < 1. According to this term, the production of *BRAVO* decreases to αεB < α when *BRAVO* is very high (*B*>> 1/K_B_) and there is no *WOX5*. The WOX5 promoter activity *P_W_* has: i) a basal production in the absence of *BRAVO* and *WOX5* expressions of value 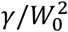; ii) *WOX5* expression ultimately represses its own production. iii) Part of this self-repression is dependent on *BRAVO*, which reduces the strength of *WOX5* self-repression. iv) The parameters *W*_0_, *B*_0_ and *W*_1_ set a measure of the characteristic *WOX5* and *BRAVO* expressions for which these regulations can have an effect.

### Modeling of the mutants

To model the mutants we used the same equations and parameter values as for the WT with the only changes being: in the *M* background (*M* can be either *bravo*, *wox5* or *bravo wox5*) the expression of the mutated gene is null at all times (*M*=0), despite its promoter activity *P_M_* is nonzero, and is computed according to the promoter function *P_M_* as defined for the WT but with *M*=0. No additional changes (e.g. no changes in parameter values) were considered to occur in the mutants. The model equations for all the mutants are detailed in Supp. Text. Herein we exemplify only the model for the *bravo* mutant (where the superscript *Bm* is used to denote this mutant):

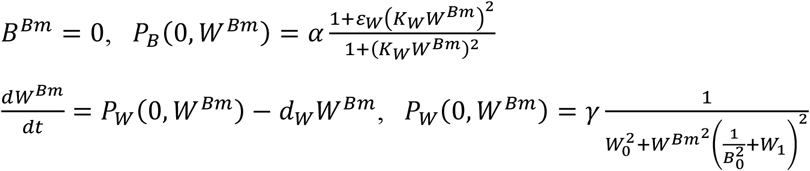

To compare with empirical data on GFP expression in the mutants, we only considered the stationary state of the mutants models (see detail in Supp. Info Text).

### Comparison of model outputs with empirical data on GFP expression

Model outputs of the promoter activities (production terms), *P_B_* and *P_W_*, obtained at the stationary state (i.e. when time-derivatives are equal to zero) were those used for comparison with the GFP data measured in the whole SCN. The superindexes *WT*, *Bm, Wm* and *dm* were used to refer to the promoter in the stationary state for the WT, the *bravo* mutant, the *wox5* mutant and the double mutant, respectively (Supp. Info Text). Since GFP scale is arbitrary with respect to promoter activity, we used the ratios that set the fold-change between mutant and the WT as the relevant measure to be compared between model outputs and empirical data. For the empirical data we used the median GFP measured values and computed the ratio of the median GFP expression in the mutant over the median GFP expression data in the WT, for each mutant. For the model, we computed the ratios of the stationary production in each mutant over its stationary production value in the WT:

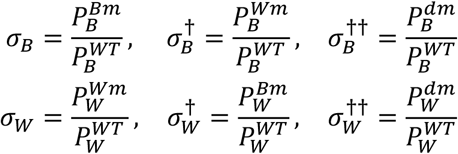

where the subscript in *σ* indicates the promoter that is analyzed (whether it is that of BRAVO or WOX5) and the superscript is informative on the mutant: no superscript is used when the ratio is evaluated in the background of the gene whose promoter is studied; superscript † is used when the mutation is on a different gene than the one driven by the promoter; †† indicates the double mutant. Parameter values in Eq.1 (Table S1) were chosen such that the values of these ratios obtained from the model fit the ratios computed from the median GFP expression values (Figure 3C). Since the GFP data is a broad distribution, there is a broad range of parameters in which the model fits the experiments within the range of experimental deviations. In addition, the model reproduces for a wide range of parameter values whether these ratios are >1 (i.e. in the mutant, the promoter activity increases) or <1 (i.e. in the mutant, the promoter activity decreases).

Notice that based on the model equations, the following equality is found for the model outputs 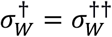 (since regulation of *WOX5* by *BRAVO* is set through *WOX5*). For *BRAVO*, 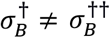 since *BRAVO* is set to self-repress, although in the range of parameters chosen both ratios are rather similar.

Additionally, the model outputs were numerically computed for different values of α and γ (all the remaining parameter values being unchanged), to model different conditions of the growth medium. Specifically, we set α and γ as functions of an auxiliary control parameter *x* that indicates the medium condition (*x*=1 corresponds to CTL conditions, whereas higher *x* values correspond to a medium with BL). We used α=0.3/*x* and γ=250*x*/(*x*+9), such that for *x*=1 α and γ take the values of the WT in CTL conditions (for *x*=1, α and γ take the values in Table S1). Roughly, *x* controls the disparity between the basal production of *BRAVO* and *WOX5*. This allows us to interpret high values of *x* as the effect of BL.

### Numerical methods to obtain model outputs

In the stationary state (i.e. when time-derivatives are equal to zero), the model for the WT reduces to a system of two coupled algebraic equations and for each mutant to a single algebraic equation (see Supp. Text). To find the stationary stable solutions we solved these algebraic equations numerically with custom-made software and using the fsolve routine embedded in Python (Python Software Foundation, https://www.python.org/), which uses a modification of Powell’s hybrid method for finding zeros of a system of nonlinear equations. The temporal evolution in Figure 3B was computed using odeint function embedded in Python (Python Software Foundation, https://www.python.org/) for the WT and for each mutant.

### Estimation of the error in the QC division data

We denote by *a,b,c* and *d* the values that we obtain empirically for the percentage of roots that exhibit a divided QC in the WT, the *bravo* mutant, the *wox5* mutant and the double *bravo wox5* mutant respectively (*a*=0.3939, *b*=0.8732, *c*=0.8070, *d*=0.8846). We can estimate the error in each of these measures, by assuming our measurement for each genotype corresponds to *N* independent equivalent roots where we observe whether the QC exhibits any division or not (i.e. we have *N* independent Bernouilli experiments). By assuming that the probability of observing a QC with at least one cell divided is *p*(*p=a,b,c,d* for each of the genotypes under study) we can estimate the error. Specifically, we assumed *p = N_k_/N*, where *N_k_* is the number of roots, from the total *N* of the specific genotype, that have a divided QC and set the error as the standard deviation of 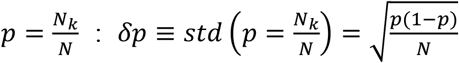. For each genotype we took a conservative view and used *N*=15 for computing the errors, so as to avoid their underestimation.

### A model to compute the contribution of BRAVO and WOX5 to regulate QC division

We aim at evaluating the contribution of BRAVO and WOX5 on regulating QC divisions. To this end we propose the following function:

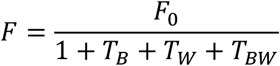

which indicates the frequency at which we found a QC with at least one QC cell that is divided in the plane of observation, for roots of the same genotype. This function can be applied to the WT, to each single mutant and to the double mutant. *T_B_*, *T_W_* and *T_BW_* are the contributions mediated by BRAVO, by WOX5 and jointly by both BRAVO and WOX5, on the regulation of QC division, such that in the *wox5* mutant we have *T_W_ =* 0 and *T_BW_* = 0, while in the *bravo* mutant we have *T_B_* = 0 and *T_BW_* = 0. Notice that for each of these contributions, it corresponds to repression of QC divisions when it takes positive values. In contrast, it corresponds to induction of QC divisions for negative values. This function takes the following expressions in the WT and in the mutants:

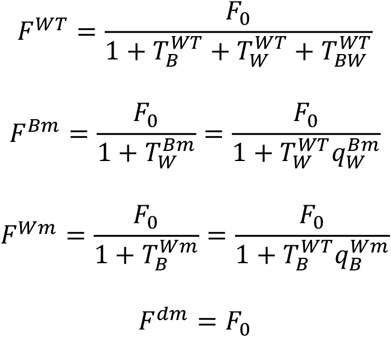

where superindexes WT, Bm, Wm account for WT, *bravo* mutant and *wox5* mutant, respectively.

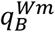 parameter measures the change in the strength of the contribution of BRAVO-mediated effects on QC division in the *wox5* mutant compared to its strength in the WT (i.e. the strength with which BRAVO inhibits QC division in the *wox5* mutant is 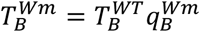). Analogously, 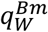 parameter measures the change in the strength of the repression that WOX5 does on QC division in the *bravo* mutant compared to the strength it does on the WT. Notice that we assume no additional changes happen in the *F* function in these mutants.

From these equations and using the empirical data (*F^WT^ = a, F^Bm^ = b, F^Wm^ = c, F^dm^ = d*, we can extract the values of 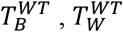 and 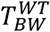 by first writing down the ratios between these quantities:

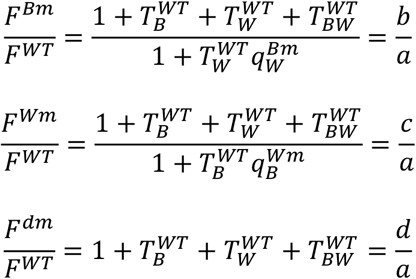

and then isolating each term, such that the following is found:

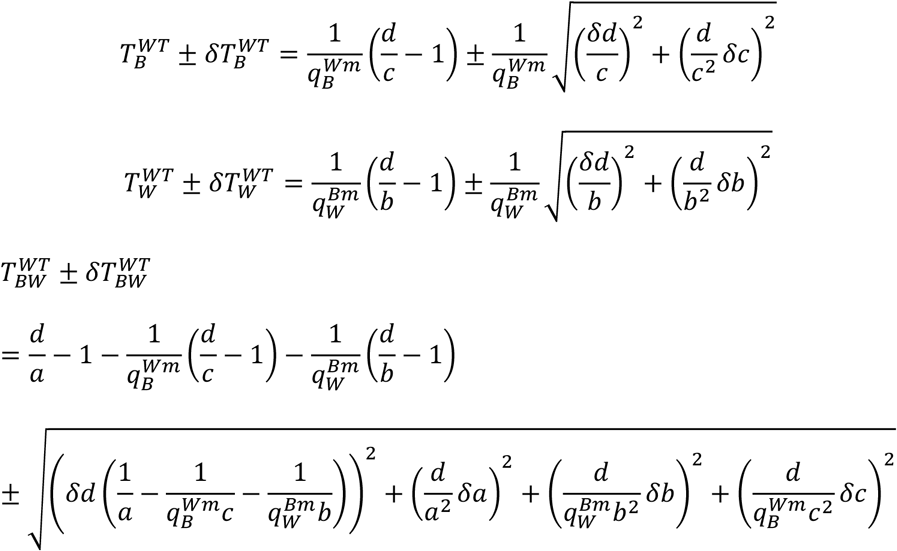

where the errors had been estimated using error propagation of the errors in *a,b,c* and *d* and assuming their independency. In Figure 5, continuous lines correspond to the best estimated values (e.g. 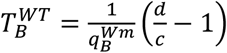), and the shaded area represents the range within the errors (e.g. 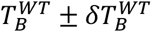). Although effective parameters 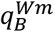 and 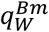 cannot be directly measured, we reasoned from the comparison of the phenotypes of *bravo* and of *wox5* mutants that 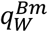 should be relatively large. As an estimate for its exact value, we used the fold-change of *WOX5* expression in the *bravo* mutant compared to the WT and set 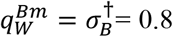. We then explored all possible values of 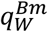 from 0 (the contribution of WOX5 in repressing divisions is eliminated completely in *bravo* mutant) to 1 (the contribution of WOX5 is the same in *bravo* mutant and in WT).

## SUPPLEMENTARY FIGURES AND TABLES

**Figure S1**: **Medial longitudinal view of the *Arabidopsis thaliana* primary root apex.**

Schematic representation of a 6-day-old primary root. At the root apex the stem cell niche is formed by the quiescent center (QC) and the surrounding stem cells, which are highlighted in different colors.

**Figure S2**: **BRAVO and WOX5 promote primary root growth and lateral root development.**

**A)** Root length of 6-day-old WT and *bravo-2 wox5-1* mutants in control and after BL treatment (n>30, 3 replicates). Different letters indicate statistically significant differences (p-value < 0.05 Student’s t-test).

**B)** Lateral root density (number of lateral roots per mm of root length) of 10-day-olf WT, *bravo-2, wox5-1* and *bravo-2 wox5-1* mutants (n>52, 3 replicates). Different letters indicate statistically significant differences (p-value < 0.05 Student’s t-test).

**Figure S3: BRAVO and WOX5 expression patterns in overexpressor lines.**

**A)** Bars show the relative expression of BRAVO and WOX5 in 35S:WOX5-GR lines when induced with 1μM Dexamethasone for 24 hours. Values in control conditions are not represented as are 1. Data obtained from two independent biological replicates. Asterisks indicate significant differences (* p-value < 0.05, *** p-value < 0.001 Student’s t-test).

**B)** Bars show the relative expression of BRAVO and WOX5 in 35S:BRAVO-Ei lines when induced with 30 μM β-estradiol for 24 hours. Values in control conditions are not represented as are 1. Data obtained from three independent biological replicates. Asterisks indicate significant differences (** p-value < 0.01 Student’s t-test).

**C)** Quantification of the GFP fluorescent signal of the roots in D-G. Boxplot indicating the average pixel intensity of the GFP in the stem cell niche. (n>29, 3 biological replicates, Different letters indicate statistical significant differences (p-value < 0.05 Student’s t-test).

**D-G)** Confocal images of PI-stained 6-day-old roots. GFP-tagged expression is shown in green. pWOX5:GFP in WT and 35S:BRAVO-Ei background in control (D, F) and after 6 days 30 μM β-estradiol induction (E, G). Scale bar: 50 μm.

**Figure S4: BRAVO expression in the *bravo wox5* mutant background.**

**A-D)** Confocal images of PI-stained 6-day-old roots. GFP-tagged expression is shown in green. *pBRAVO:GFP* in WT and *bravo-2 wox5-1* background in control (A, C) and after BL treatment (B, D). Scale bar: 50 μm.

**E)** Quantification of the GFP fluorescent signal of the roots in A-D in the stem cell niche. Different letters indicate statistically significant differences (p-value < 0.05 Student’s t-test).

**Figure S5: BRAVO and WOX5 expression is BL regulated.**

**A-N)** Confocal images of PI-stained 6-day-old roots. GFP-tagged expression is shown in green. **A-C)** *pBRAVO:GFP* in WT, *bravo-2* and *wox5-1* knockout backgrounds in CTL (A-C) and after 48h 4nM BL treatment (D-F). **G-N)** *pWOX5:GFP* in WT, *bravo-2, wox5-1* and *bravo-2 wox5-1* knockout backgrounds in CTL (G-J) and after 48h 4 nM BL treatment (K-N). Images in control conditions are the same that are shown in figure 2. Scale bar: 50 μm.

**O, P)** Quantification of the GFP fluorescent signal of the roots in A-F (O) and G-N (P). Boxplot indicating the average pixel intensity of the GFP in the stem cell niche. (n>25, 3 biological replicates, *p-value < 0.05 Student’s *t*-test for each genotype versus the WT in the same condition). Quantification of lines in control conditions are the same that are shown in figure 2.

**Figure S6: Biochemical interactions of BRAVO and WOX5 with BES1 and TPL.**

**A)** Yeast two-hybrid assay showing BRAVO interactions with WOX5, BES1 and TPL *in vitro*. In the left column yeast cells were grown on control media, and in the right column yeast cells were grown on control media lacking Leu, Trp and His, indicating an interaction between the proteins.

**B-D)** *In planta* interaction by Bimolecular Fluorescence Complementation assay (BiFC). Confocal images were merged with red fluorescence images corresponding to chlorophyll. Fluorescence was detected 48 h post agroinfiltration. Scale bar: 50 μm. B) BiFC showing BRAVO interaction with BES1 and TPL. Nuclear YFP fluorescence is observed in *N. benthamiana* leaves infiltrated with the BRAVO-eYFPC and both BES1 and TPL-eYFPN constructs. BRAVO-eYFPC and empty-eYFPN are included as a negative control. C) BiFC showing WOX5 interaction with BES1 and TPL. Nuclear YFP fluorescence is observed in *N. benthamiana* leaves infiltrated with the WOX5-eYFPN and both BES1 and TPL-eYFPC constructs. WOX5-eYFPN and empty-eYFPC are included as a negative control. D) BES1-eYFPC and TPL-eYFPN was included as a positive control of interaction. Scale bar: 50 μm.

**Figure S7: ROIs used for the quantification of the GFP.**

**A-B)** Confocal images of *pBRAVO:GFP* (A) *and pWOX5:GFP* (B) PI-stained 6-day-old roots. GFP-tagged expression is shown in green. Insets show the GFP channels that were used for the quantification. Only the area inside the yellow circle was used for the GFP quantification.

**Table S1. Parameter values for the model of BRAVO and WOX5, used to generate the data in Figure 3.**

Parameter values used to perform the numerical simulations. All are in arbitrary units. The right-most column indicates the concentration and time scales in which these values could be meaningful in a biological context.

**Table S2. List of plant material lines used in this study.**

**Table S3. List of primers used in this study.**

## SUPPLEMENTARY TEXT

### Model

For the WT genotype, the model reads (see Material and Methods):

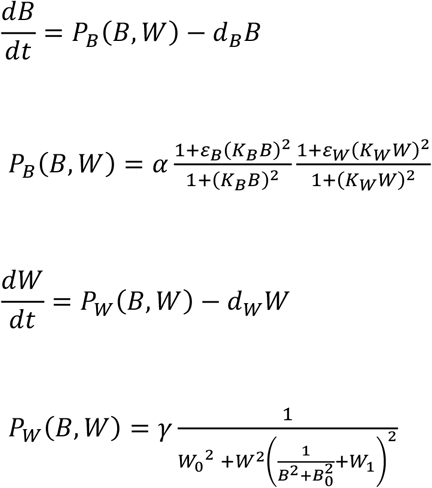

For the *wox5* mutant (where superscript *Wm* denotes this mutant) the model reads (it has *W^Wm^* = 0):

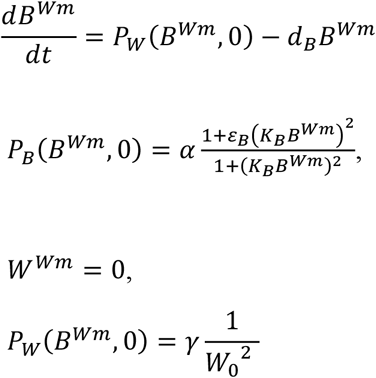

The model for *bravo* mutant (where superscript *Bm* denotes this mutant) has *B^Bm^ =* 0 and reads:

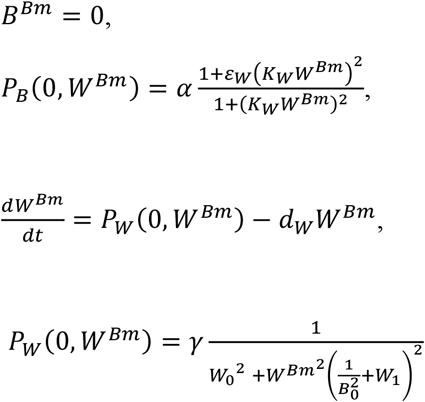

Finally, for the double *bravo wox5* mutant (superscript *dm*) the model reads:

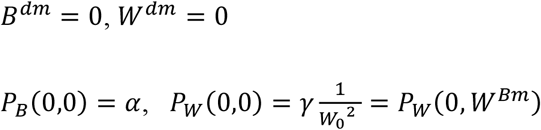

### Stationary solutions

For each genotype, the stationary solutions are found by imposing the stationarity condition: 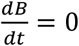 and 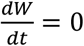, of the equations that describe each genotype.

For the WT, when we impose the stationary conditions the following set of two coupled algebraic equations is obtained in the stationary state:

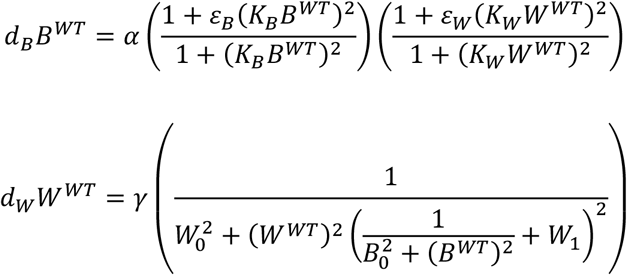

which is solved numerically (see Material and Methods). We denote by *B^WT^, W^WT^* the stationary solutions for the expression of *BRAVO* and *WOX5* in the WT. The stationary *BRAVO* and *WOX5* promoter activities in the WT are:

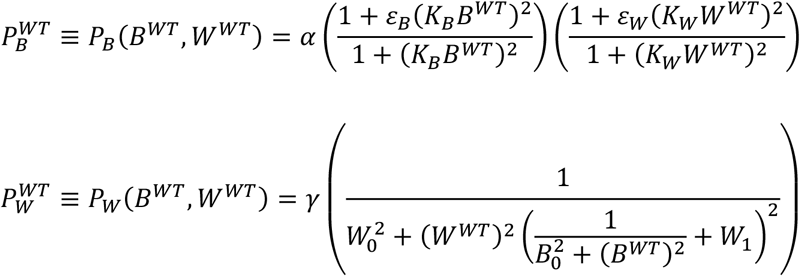

where, once we have the stationary values *B^WT^, W^WT^* we can obtain their values by substitution on the above expressions.

We proceed in the same way with each mutant with their corresponding equations set to the stationary state.

For the *wox5* mutant, we have *W^Wm^* = 0, and the stationary expression of *BRAVO* satisfies

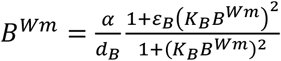

which is solved numerically. The stationary *BRAVO* and *WOX5* promoter activities (productions) in this mutant are:

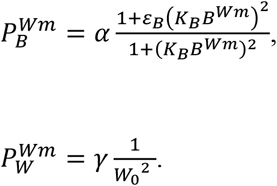

For the *bravo* mutant in the stationary state we have *B^Bm^* = 0, and

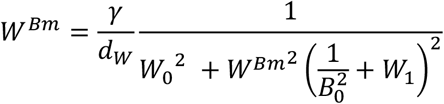

which is solved numerically. Once solved, the stationary promoter activities in this mutant are found as:

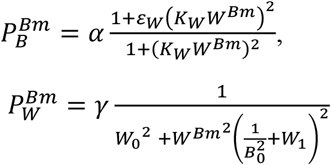

Finally, the model of the double *bravo wox5* mutant already indicates the stationary state values:

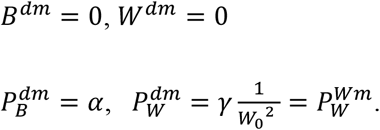

